# Parental effects can be transmitted through the rhizosphere microbiome

**DOI:** 10.64898/2026.11.12.766069

**Authors:** Yu-Han Chen, Jun Hee Jung, Frank Reis, Madalin Parepa, Chun-Lin Wang, Tao Fang, Yi-Fan Liu, Jian-Qiao Meng, Fang-Li Luo, Oliver Bossdorf

## Abstract

- Plant responses to environmental changes, including to competition by neighboring plants, can be influenced by soil microbes. They can also be modified through parental effects, which can transmit environmental cues across generations and shape plant phenotypes through various mechanisms such as epigenetic modifications. However, we do not yet know whether parental effects in plants can also act through modifications of the rhizosphere microbiota.
- Here, we grew eight genotypes of *Phragmites australis* with or without interspecific competition, and examined its transgenerational effects on offspring phenotype and rhizosphere microbial communities in a common environment.
- We found that competition decreased plant growth in a genotype-specific manner, but plants whose parents had been exposed to competition previously shifted their functional phenotypes towards a more conservative resource strategy. Notably, parental competition also altered the diversity and community structure of rhizosphere soil bacteria and fungi in offspring plants, and structural equation models indicated that these soil microbiome changes may have driven some of the observed changes in offspring phenotypes.
- Our study demonstrates that parental environmental effects can influence offspring traits not only directly, but also through alteration of rhizosphere microorganisms, and that parental effects are more complex than we previously knew.

## Introduction

Plants are sessile organisms, and they must therefore be able to adjust to changing habitat conditions, sometimes very rapidly. So far, the main mechanism for explaining such rapid adjustments has been phenotypic plasticity - a suite of developmental and regulatory mechanisms that alter plant phenotypes in response to environmental changes (Gratani, 2014; Palacio-López *et al*., 2015; Sammarco *et al*., 2022). However, there is increasing evidence that plant-associated microorganisms, too, may play a role in rapid plant adaptation (Rezki *et al*., 2018; Bai *et al*., 2022). In particular, below-ground microbial communities can influence plant growth and physiology, e.g. through altering nutrient use efficiency, synthesizing phytohormones and secondary compounds, and thereby modulating immune responses and plant defenses, or inhibiting pathogens (Li *et al*., 2021b; Geisen *et al*., 2022; Wang *et al*., 2024). These processes can not only affect the stress tolerance of individual plants, but also the structure and diversity of plant communities (Das *et al*., 2022; Zieschank *et al*., 2025). In competitive environments, microbes can influence plant resource use and nutrient uptake, and specific microbiota may determine the outcome of interspecific plant competition (Hodge & Fitter, 2013). While the impact of specific microbes on plant growth can be positive, neutral or negative (Revillini *et al*., 2016; Aqeel *et al*., 2023), a higher diversity of rhizosphere microbial communities is generally expected to have positive effects on plant growth and stress resistance (Lau & Lennon, 2011). Finally, plant-microbe interactions are generally reciprocal. The physiological states of the plants can feed back on the microbes e.g. through root exudates, and therefore the stress exposure of plants, but also their age or genotype, can influence plant-associated microbial communities (Lareen *et al*., 2016; Wagner *et al*., 2016; Pattnaik *et al*., 2021; Shelake *et al*., 2026).

Besides the direct effects of the environment, plant phenotypes can also be influenced by the environments experienced by their parents, a phenomenon known as ‘parental effects’ or ‘transgenerational plasticity’ (Yin *et al*., 2019; McAndry *et al*., 2025). Parental effects occur both in sexually reproducing and clonal plants (Latzel *et al*., 2024; Sammarco *et al*., 2025), i.e. through meiotic or mitotic ‘memory’. In both cases, physiological signals caused by particular environmental stresses such as drought, cold, heat, salt or light are transmitted to offspring plants through a variety of mechanisms such as nutrient provisioning, epigenetic modifications, etc., with significant effects on their phenotypes (Miryeganeh & Saze, 2020; Latzel *et al*., 2024). Transgenerational effects can be adaptive, i.e. increase the fitness of offspring, but they can sometimes also be detrimental (Dong *et al*., 2018). One aspect of transgenerational effects that we so far know nothing about is the role of plant-associated microbes, which could be influenced transgenerationally, but they may also be mediators of transgenerational effects if they feed back on the plant phenotypes.

In nature, plants do not grow in isolation, but they usually interact with competitors. Interspecific competition can alter resource availability and soil conditions (Craine & Dybzinski, 2013; Springer *et al*., 2024), thereby affecting plant reproductive strategy and offspring phenotypes transgenerationally (Germain *et al*., 2019; Waterman & Sultan, 2021). For instance, in the perennial apomictic herb *Taraxacum brevicorniculatum*, parental competition reduced the leaf decomposability of offspring plants, and resulted in more resource-conservative phenotypes (Puy *et al*., 2021). However, there are still only few studies on parental effects caused by interspecific competition, especially for clonal plants (Yu *et al*., 2022; He *et al*., 2025).

The phenotypes of plants and their biotic interactions are usually partly genotype-dependent (Hughes *et al*., 2008; Andrade *et al*., 2019), and parental effects, too, can vary considerably among genotypes (Ibañez *et al*., 2017; Deng *et al*., 2021; Zhang *et al*., 2025). Therefore, a robust general understanding of transgenerational plasticity in plants requires to study such effects across multiple genotypes (Münzbergová *et al*., 2017). Clonal plants are excellent study systems for addressing such questions, not only because their predominantly vegetative reproduction allows a broader transmission of parental memories than if genetic information is reset through meiosis (Verhoeven & Preite, 2014), but also because of the easy production of genetically identical replicates of different plant genotypes for experimental comparisons.

Here, we worked with *Phragmites australis* (Cav.) Trin. ex Steud., a perennial grass that is widely distributed in wetlands around the world. It is a typical clonal plant with a high capacity for clonal reproduction, but it is also capable of sexual reproduction. Presumably because of its broad natural distribution and long-term adaptation to contrasting environmental conditions, such as salinity and flooding, *P. australis* is genetically very diverse (Fant *et al*., 2016; Wang *et al*., 2021). The *P. australis* populations in salt marshes along the east coast of China play an important role in maintaining the structure and function of coastal wetland ecosystems (Jia *et al*., 2024). However, in these environments, *P. australis* now co-occurs with the invasive *Spartina alterniflora*, a strong competitor with efficient nutrient utilization and photosynthesis, and high clonal reproduction (Shang *et al*., 2015; Liu *et al*., 2023a). *S. alterniflora* is currently spreading rapidly, gradually limiting the growth of native plants such as *P. australis*, and threatening the stability of coastal wetland ecosystems (Meng *et al*., 2020). It is an open question whether in the long term the native *P. australis* will be able to adapt to the competitive pressure from *S. alterniflora*, and it is therefore important to explore potential parental effects of *S. alterniflora* competition, as well as the variation in competitive ability and parental effects among different genotypes of *P. australis*.

Below, we present a two-generation common-garden experiment (Fig. 1) in which we tested for transgenerational effects of interspecific competition by *S. alterniflora* on the phenotypes and rhizosphere microbial communities of *P. australis*. We worked with eight natural genotypes of *P. australis*, collected from four estuarine wetlands along the east coast of China, and we addressed the following questions: (1) How does competition in the parental generation affect the phenotypes of offspring plants, and if so, is it genotype-dependent? (2) Do parental competition and parental traits influence the diversity and structure of rhizosphere bacteria and fungi associated with offspring plants? (3) Are transgenerational changes in rhizosphere microbes related to changes in offspring phenotypes?

**Fig. 1.**
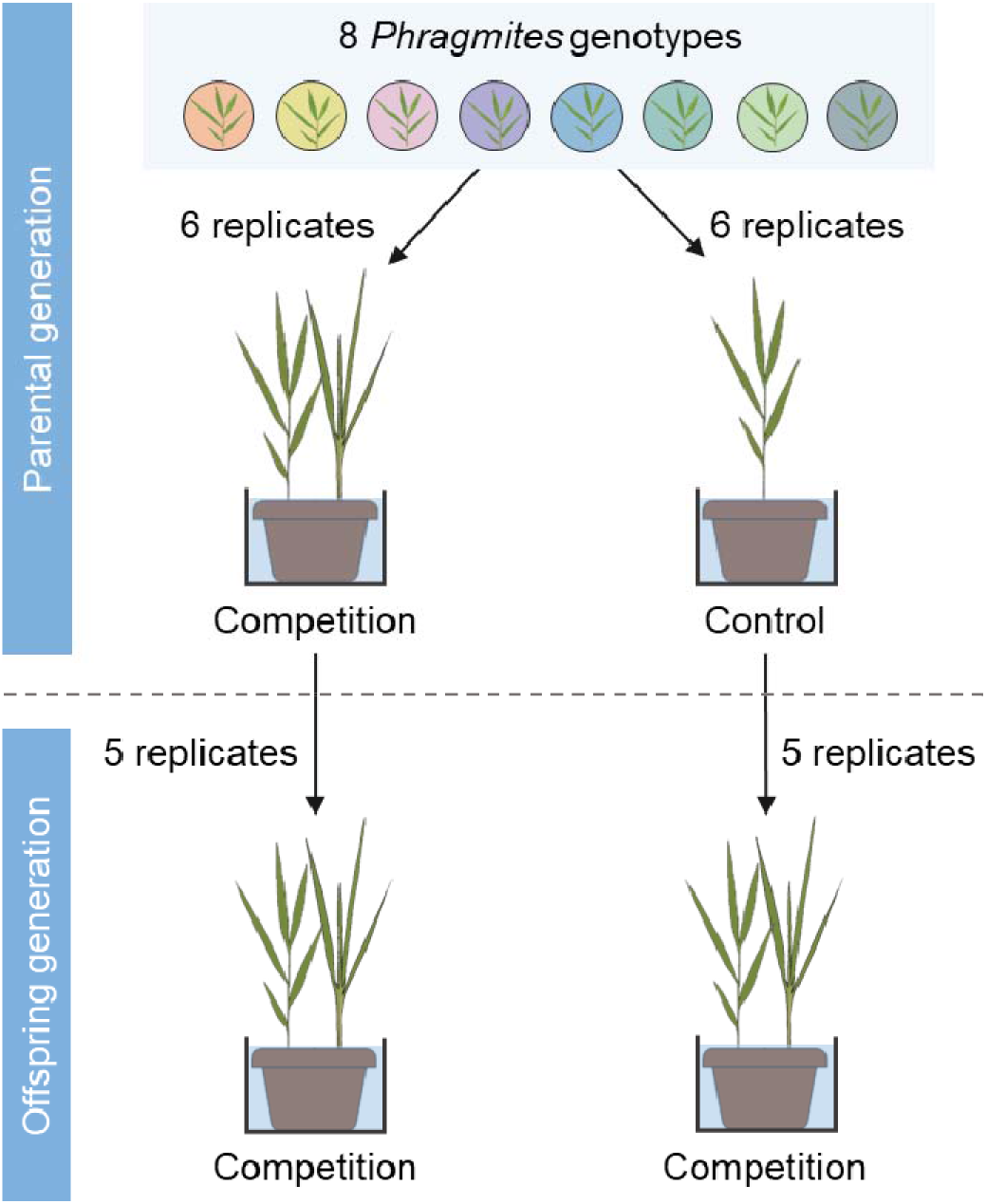
Schematic of the design of our two-generation competition experiment, with eight genotypes of *Phragmites australis* collected from four estuarine wetlands along the Chinese eastern coast, and 12 seedlings per genotype grown with or without interspecific competition by *Spartina alterniflora* in the parental generation, and then their offspring grown under competition in the second generation.

## Material and Methods

### Plant collection

We worked with eight genotypes of *P. australis* that we selected from a larger survey of 42 natural populations across four estuarine wetlands in China. The estuaries covered a large latitudinal gradient, and in each the populations were evenly distributed. To represent the natural variation within this species, we selected, based on a phylogenetic tree constructed from Simple Sequence Repeat marker data (Fig. S1), eight genotype with relatively large genetic distances (Chen *et al*., 2024). We collected rhizomes between October and November 2020 and pre-cultivated all plants in a common greenhouse environment at Beijing SFK Technology Co., Ltd., China (40°0′27"N, 116°20′19"E) for 10 months to minimize differences induced by the habitat of origin.

### Experimental set-up

To investigate parental effects of interspecific competition, we conducted an experiment over two generations (Fig. 1). In the parental generation, we set up a competition experiment in which we used *S. alterniflora* plants, collected close to Yancheng (32°46′56"N, 120°57′42"E) in the Jiangsu Province, as competitors. For each *P. australis* genotype we planted 12 uniformly sized seedlings, each with a 10 cm rhizome, individually into separate pots filled with a 1:1 mixture of sand and loam. In half of the pots of each genotype, we added one *S. alterniflora* plant per pot as a competitor, always at the same position and distance to *P. australis*. We further added a 10‰ sea salt solution to all pots, maintaining the water level 1 cm above the soil surface, to mimic the moderate salinity of the estuary environment where both species coexist (Li *et al*., 2020). There were six replicates per genotype and treatment, and a total of 96 pots (eight genotypes x two treatments x six replicates). We placed the pots in a greenhouse, in a randomized block design with six blocks and each block containing one plant from each genotype and treatment. After four months, in December 2021, we harvested all plants, measured a range of phenotypic and physiological traits (see below), and then cut and cleaned all 10 cm rhizome tips of each plant and measured their fresh weights. We used the newest 10 cm rhizome tips of five replicates per genotype to establish the offspring generation, this time all with competition, i.e. *P. australis* and *S. alterniflora* planted together in each pot. We used the same pots and substrate as in the parental generation. The offspring experiment consisted of 80 plants (eight genotypes x two parental treatments x five replicates), distributed across five randomized blocks, and it lasted from June to October 2022. Unfortunately, some rhizomes did not resprout, and at the end of the experiment we harvested only 53 plants.

### Phenotype measurements

At the end of the parental and offspring experiment, we collected nearly identical phenotype data. We measured the height and basal stem diameter of the tallest *P. australis* ramet in each pot, as well as the leaf area of its largest fully expanded mature leaf (hereafter maximum leaf area). We then carefully cleaned the belowground parts of each *P. australis* plant, counted the number of its rhizome nodes as an estimate of clonal reproduction, and determined the combined fresh weight of all 10 cm rhizome tips, as well as the fresh weight of the remaining belowground parts. We separated the second spacer (rhizome section between second and third nodes from the base) of the tallest ramet and then determined its dry biomass, as well as that of all above- and below-ground plant parts, after drying at 70°C for 72 h. We used the moisture content derived from the remaining belowground parts to estimate the dry weight of the rhizome tips for further total dry biomass calculation. We calculated growth allocation as aboveground dry biomass divided by total dry biomass. Prior to the harvest of the offspring plants, we additionally characterized their photosynthetic capacity through measuring the quantum yield of photochemical energy conversion in PS II (Y(II)), using a PAM-2500 chlorophyll fluorometer (Heinrich Walz GmbH, Germany).

### Microbiome sequencing

Prior to harvesting the *P. australis* offspring plants, we collected rhizosphere soil around their roots, removed the fine roots through a 2 mm sieve, and immediately froze the samples at −80°C. For each genotype and parental treatment, we collected soil from three replicate plants, for a total of 48 soil samples. To characterize the bacterial and fungal microbial communities in the rhizosphere soil samples, we employed 16S and ITS2 amplicon sequencing. DNA extraction and library preparation, as well as DNA sequencing, were done at Novogene (Beijing, China), following their standardized protocols for soil sample processing and sequencing. Briefly, after extraction of total genomic DNA, and control of DNA concentration and purity, the V4 regions of the 16S rRNA genes were amplified using primers 515F (5′-GTGCCAGCMGCCGCGGTAA-3′) and 806R (5′-GGACTACHVGGGTWTCTAAT-3′), and the internal transcribed spacer (ITS2) regions were amplified using primers ITS3-2024F (5′-GCATCGATGAAGAACGCAGC-3′) and ITS4-2409R (5′-TCCTCCGCTTATTGATATGC-3′). Each PCR reaction included 15 μL of Phusion High-Fidelity PCR Master Mix (New England Biolabs), 0.2 μM of each primer, and 10 ng target DNA, and consisted of the following steps: initial denaturation at 98°C for 1 min, 30 cycles at 98°C for 10 s, 50°C for 30 s, 72°C for 30 s, and final extension at 72°C for 5 min. The PCR products were detected through 2% agarose gel electrophoresis, mixed in equal proportions, purified using the Qiagen Gel Extraction Kit (Qiagen, Germany), and sequenced on the Illumina NovaSeq 6000 platform to generate 250 bp paired-end reads. After removing barcodes and primer sequences, the paired-end reads were then truncated by cutting off barcodes and primer sequences. We used DADA2 (Callahan *et al*., 2016) and QIIME2 version 2024.5 (Bolyen *et al*., 2019) for denoising, chimera checking and joining of the truncated paired-end reads. The truncation and trimming were set to --p-trim-left-f 0, --p-trim-left-r 0 and --p-trunc-len-f 200, --p-trunc-len-r 200 for the 16S and ITS data. We then used QIIME phylogeny align-to-tree-mafft-fasttree to build a phylogenetic tree based on the filtered reads, and we applied the QIIME feature-classifier classify-sklearn with the Silva 16S v4 database (Silva 138 SSURef NR99 515F/806R region sequences, https://docs.qiime2.org/2024.10/data-resources/) for the taxonomic assignment of 16S data (Quast *et al*., 2013; Yilmaz *et al*., 2014; Glöckner *et al*., 2017) and with the UNITE v10.0 reference datasets (sh_refs_qiime_ver10_99_04.04.2024_dev, https://doi.plutof.ut.ee/doi/10.15156/BIO/2959336) for the taxonomic assignment of the ITS2 data (Abarenkov *et al*., 2020).

### Data analyses

All analyses were carried out using R v.4.4.2 (R Core Team, 2024). To test the direct effects of competition on traits in the parental generation, we fitted linear mixed-effect models that included plant genotype (G), competition environment (C; with or without competition) and their interaction (G × C) as fixed factors and experimental block as a random factor. To meet the assumptions of normality and homogeneity of variance of the residuals, we log-transformed the height, maximum leaf area, rhizome nodes and total biomass data prior to the analyses. Next, we examined the effects of parental competition on offspring phenotypes with linear mixed-effect models that included plant genotype (G), parental competition (C) and their interaction (G × C) as fixed factors. For all growth traits, we included initial weight as a covariate, and we generally included experimental block as a random effect. Prior to the analysis, we log-transformed the height, basal stem, rhizome nodes and total biomass data, and sqrt-transformed the maximum leaf area, spacer biomass and Y(II) data. We used the *lme4* and *lmerTest* packages to conduct these analyses (Bates *et al*., 2015; Kuznetsova *et al*., 2017), extracted the predicted values from these models using the *ggeffect* function in the *ggeffects* package (Lüdecke, 2018), and visualized the results using the *ggplot2* package (Wickham, 2016).

For the microbial data, we imported the QIIME2 result files into R and removed all sequences identified as mitochondria, chloroplast, human or archaea. We then used the *microeco* package (Liu *et al*., 2021) for the bacterial and fungal data analyses. We estimated microbial alpha diversity through the Shannon–Wiener index and analyzed these diversity data with linear mixed-effect models that had the same structure as for the offspring phenotypes. To enhance model robustness, we excluded outliers with standardized residuals exceeding 3.0. This excluded in particular one extreme outlier (G7-CP3) in bacterial alpha diversity from the subsequent analyses. To characterize the taxonomic beta diversity among different samples, we calculated a dissimilarity matrix based on Bray-Curtis distances, carried out principal coordinates analyses (PCoA) of these Bray-Curtis distances, and tested for significant group differences in bacterial or fungal community structure with permutational multivariate analysis of variance (PERMANOVA) with 99999 permutations in the *vegan* package (Anderson, 2017; Oksanen *et al*., 2025). To identify specific differentially abundant microbial taxa between different genotypes or parental competition environments, we calculated Linear Discriminant Analysis Effect Sizes (LEfSe) (Segata *et al*., 2011), using a significance threshold of α = 0.05 for the Kruskal-Wallis test. Features (= microbial taxa) that showed significant differences were further evaluated based on their effect sizes, with an LDA score threshold of 2.0 (except for bacterial taxa between different plant genotypes where we used a threshold of 3.0). Finally, we used Mantel tests to test for relationships between the (dis-)similarities of plant-associated microbial communities and plant traits, separately for bacterial and fungal communities, and for the parental and offspring generation (Liu *et al*., 2017).

To better understand direct versus indirect relationships between soil microbial diversity and plant traits in the parental and offspring generation, we employed structural equation modelling (SEM; *lavaan* package in R) (Rosseel, 2012). Prior to the SEM analyses, we standardized all variables and, when necessary, transformed them to achieve normality. We also tested for collinearity of variables using Pearson’s correlations and removed variables with *r* > 0.8), and we tested for variance inflation using the *vif* function of the *car* package (Fox & Weisberg, 2019). After excluding the redundant variables, we first created a complex initial model that included all remaining variables in a plausible causal structure, and we then removed non-significant variables to find the best SEM with the lowest AIC. We evaluated the overall model fit with a Chi-square test (*p* > 0.05), a comparative fit index (CFI > 0.95), and the root mean squared error of approximation (RMSEA < 0.08).

## Results

### Effects of competition on plant phenotypes

Interspecific competition significantly reduced the total biomass of *P. australis* plants in the parental generation (Table 1; Fig. S2E), and it affected the height, basal stem diameter, spacer biomass and growth allocation of the plants in a genotype-specific manner (significant G x C interactions in Table 1; Fig. S2). In all traits but the number of rhizomes nodes, there were significant differences among plant genotypes (Table 1).

**Table 1.** Results of analysis of variance testing the effects of genotypes (G), parental competition (C) and their interaction (G x C) on phenotypes of *Phragmites australis* in the parental and offspring generation. The values are *F*-values, with significant ones in bold: *** *P* < 0.001, * *P* < 0.05

|  | Traits | Initial weight | G | C | G × C |
|---|---|---|---|---|---|
| Parental | Height | **-** | **13.74**<sup>\*\*\*</sup> | 0.00 | **2.22**<sup>\*</sup> |
|  | Basal stem diameter | **-** | **4.46**<sup>\*\*\*</sup> | 3.48 | **2.54**<sup>\*</sup> |
|  | Maximum leaf area | **-** | **11.59**<sup>\*\*\*</sup> | 1.95 | 0.55 |
|  | Spacer biomass | **-** | **6.65**<sup>\*\*\*</sup> | 0.26 | **2.37**<sup>\*</sup> |
|  | Rhizome nodes | **-** | 1.47 | 1.70 | 1.49 |
|  | Total biomass | **-** | **5.59**<sup>\*\*\*</sup> | **6.16**<sup>\*</sup> | 0.97 |
|  | Growth allocation | **-** | **8.94**<sup>\*\*\*</sup> | 0.28 | **2.37**<sup>\*</sup> |
| Offspring | Height | 2.42 | **6.58**<sup>\*\*\*</sup> | 2.04 | 0.93 |
|  | Basal stem diameter | 2.82 | **5.80**<sup>\*\*\*</sup> | 0.17 | 1.16 |
|  | Maximum leaf area | 0.59 | 0.71 | **5.86**<sup>\*</sup> | 0.72 |
|  | Spacer biomass | **5.21**<sup>\*</sup> | 1.03 | 0.27 | 0.18 |
|  | Rhizome nodes | 0.05 | **2.91**<sup>\*</sup> | 0.002 | 0.61 |
|  | Total biomass | 2.38 | **2.44**<sup>\*\*\*</sup> | 0.36 | 0.56 |
|  | Growth allocation | 0.25 | **2.71**<sup>\*\*\*</sup> | 0.05 | 0.85 |
|  | Y(II) | **-** | **3.03**<sup>\*\*\*</sup> | 0.39 | **2.37**<sup>\*</sup> |

In the offspring generation, the parental competition environment consistently impacted the maximum leaf area (Table 1). Plants whose parents had been growing under interspecific competition had a 17.14% lower maximum leaf area than plants whose parents did not experience competition (Fig. 2A). Parental competition also influenced the photosynthetic capacity (Y(II)) of offspring plants, albeit in a genotype-specific manner (Fig. 2B). There were no further parental effects on the other traits of offspring plants. As in the parental generation, we also found significant genotype effects on most (6 out of 8) of the offspring phenotypic traits (Table 1; Figs. 2 and S3).

**Fig. 2.**
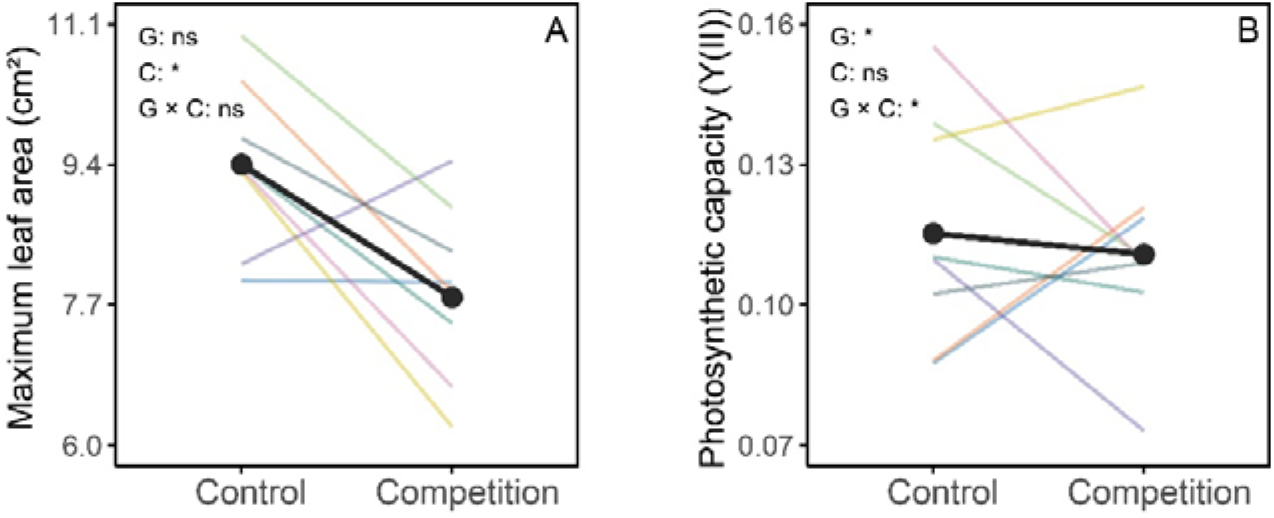
Effects of parental competition environment on the (A) maximum leaf area and (B) Y(II) (photosynthetic capacity) of offspring plants in *P. australis*. The coloured lines represent different genotypes, and the thick black line is the main effect (across genotypes) of parental competition. All values are predicted values from linear mixed-effects models, back-transformed to the original scale. The significances of fixed effects (G: genotype, C: parental competition) are indicated in the figure panels, with * *P* < 0.05 and ns *P* > 0.05.

### Diversity and structure of offspring soil microbes

The α- and β-diversity of offspring rhizosphere soil bacteria as well as the community structure of fungi varied notably among plant genotypes (Fig. 3). LEfSe analysis identified differential abundance of several bacterial families, but only few fungi taxa, among different *P. australis* genotypes (Fig. S4B). Moreover, parental competition significantly influenced the community composition of both rhizosphere soil bacteria and fungi (Fig. 3C, D). Again, LEfSe identified a large number of bacterial families significantly associated with parental competition, with GAL15 as the most enriched and *Prolixibacteraceae* as the most strongly decreased bacterial family (Fig. S4A), but only few significantly differentiated fungi taxa (Fig. S4C). We found a significant interaction between genotype and parental competition in the analysis of bacterial Shannon diversity (Fig. 3A), indicating that parental effects were strongly genotype- dependent for this aspect of the root microbiome.

**Fig. 3.**
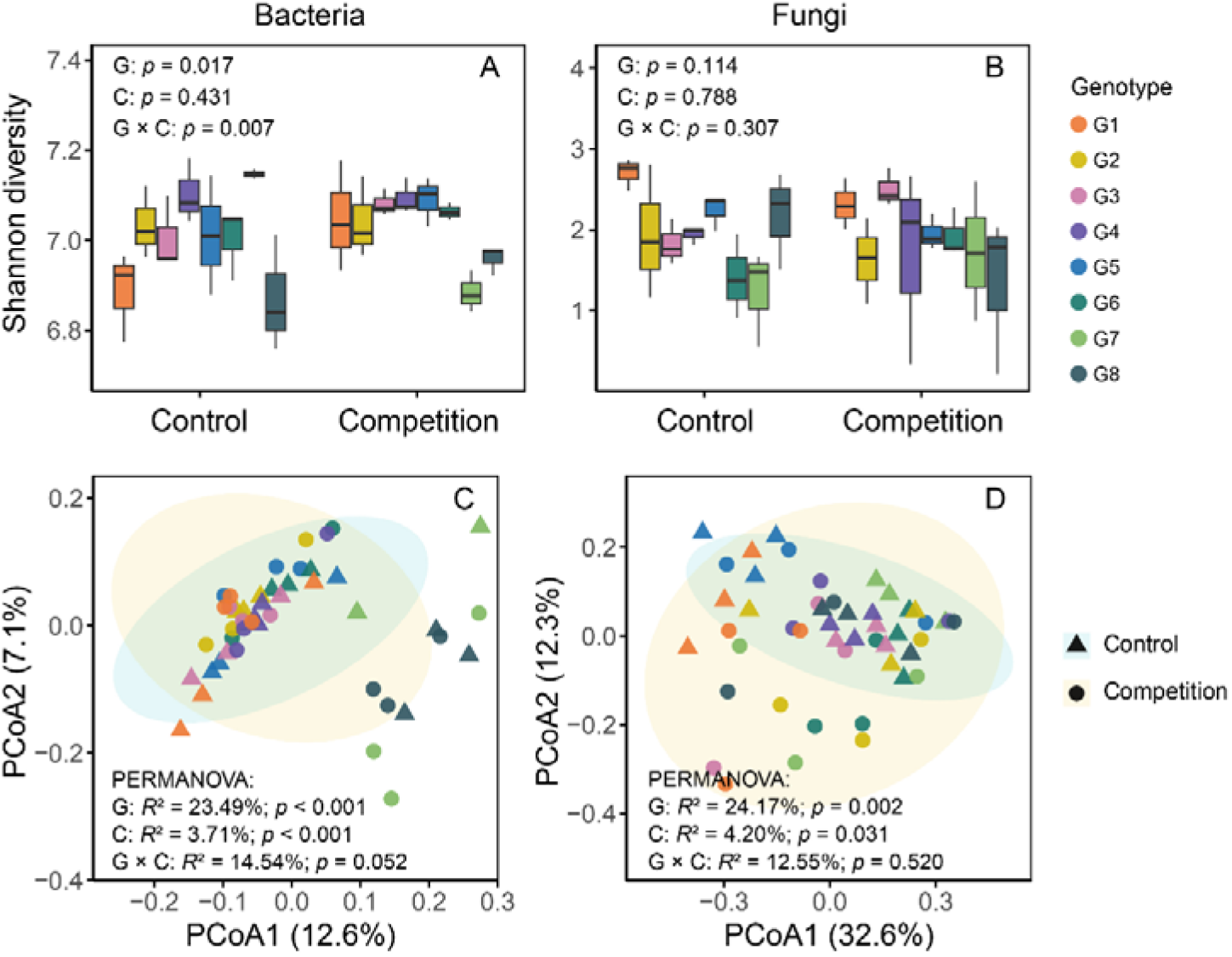
Diversity of rhizosphere soil microorganisms associated with different genotypes of offspring *P. australis* under parental control or parental competition: (A, B) Shannon diversity of bacteria and fungi; (C, D) Community (dis)similarities (PcoA = principal coordinate analysis based on Bray-Curtis distances) of bacterial and fungal communities associated with different plant genotypes and parental treatments.

### Relationships among soil microbes, parental traits and offspring traits

The observed variation in offspring rhizosphere microbial communities was associated with variation in plant phenotypes. However, parental competition altered these relationships (Fig. 4). For the offspring from parents without competition, the composition of rhizosphere bacteria was significantly associated with the stem diameter and spacer biomass of the parents, but for plants whose parents had experienced competition, the bacterial communities were instead associated with parental variation in plant height and growth allocation. Interestingly, we also found significant associations between rhizosphere microbes and offspring phenotypes, in particular between bacterial community structure and plant stem diameter, and between fungal community structure and maximum leaf area. However, these associations occurred only in offspring from plants that had previously experienced competition (Fig. 4).

**Fig. 4.**
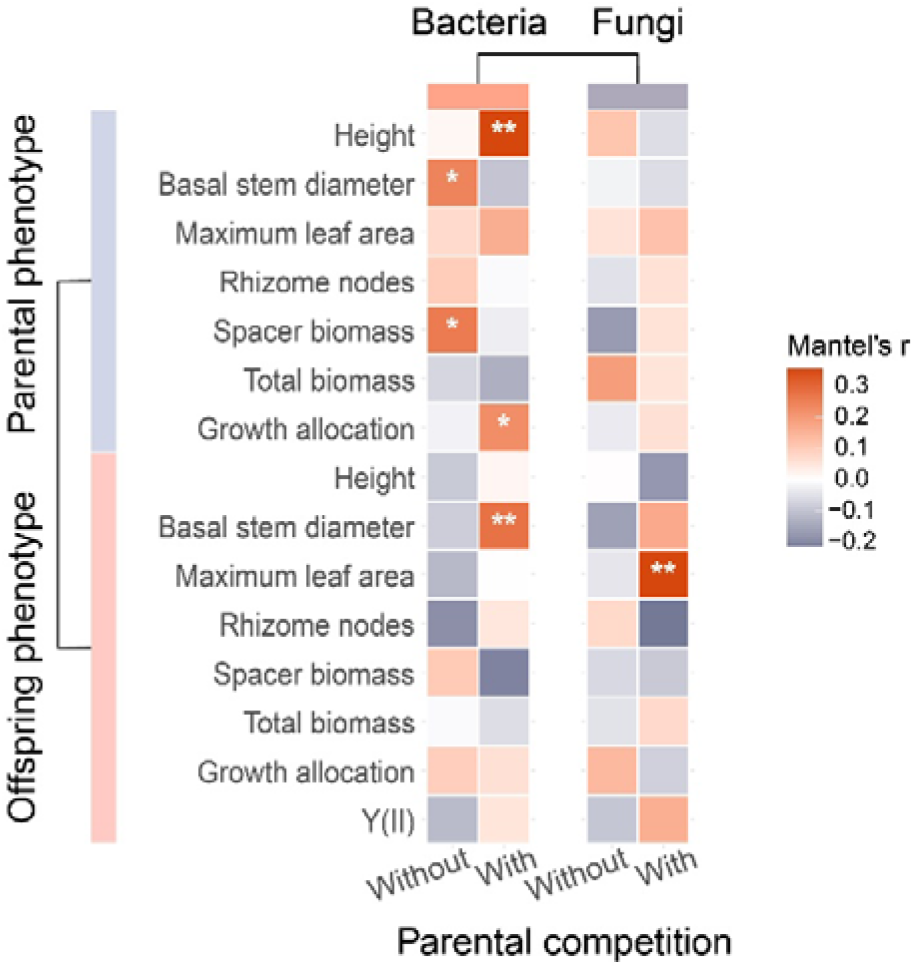
Results of Mantel tests of the relationships between parental or offspring phenotype and bacterial or fungal community composition of *P. australis*, with orange values indicating positive correlations and blue negative ones, and significance levels as **0.001 ≤ *P* < 0.01, *0.01 ≤ *P* < 0.05.

When we connected parental competition environment, parental and offspring traits, and offspring rhizosphere microbes through structural equation modeling (SEM, Fig. 5), we found significant negative paths from parental competition to both parent and offspring traits, but also a negative path from the parental traits to the offspring traits, thus adding an indirect positive effect of parental competition on the offspring traits. The final SEM also contained positive paths from parental competition and parent traits to the rhizosphere bacterial diversity of the offspring plants, which in turn had a positive effect on offspring traits (Fig. 5), thus adding multiple additional microbe-mediated pathways through which parental competition affected offspring phenotypes.

**Fig. 5.**
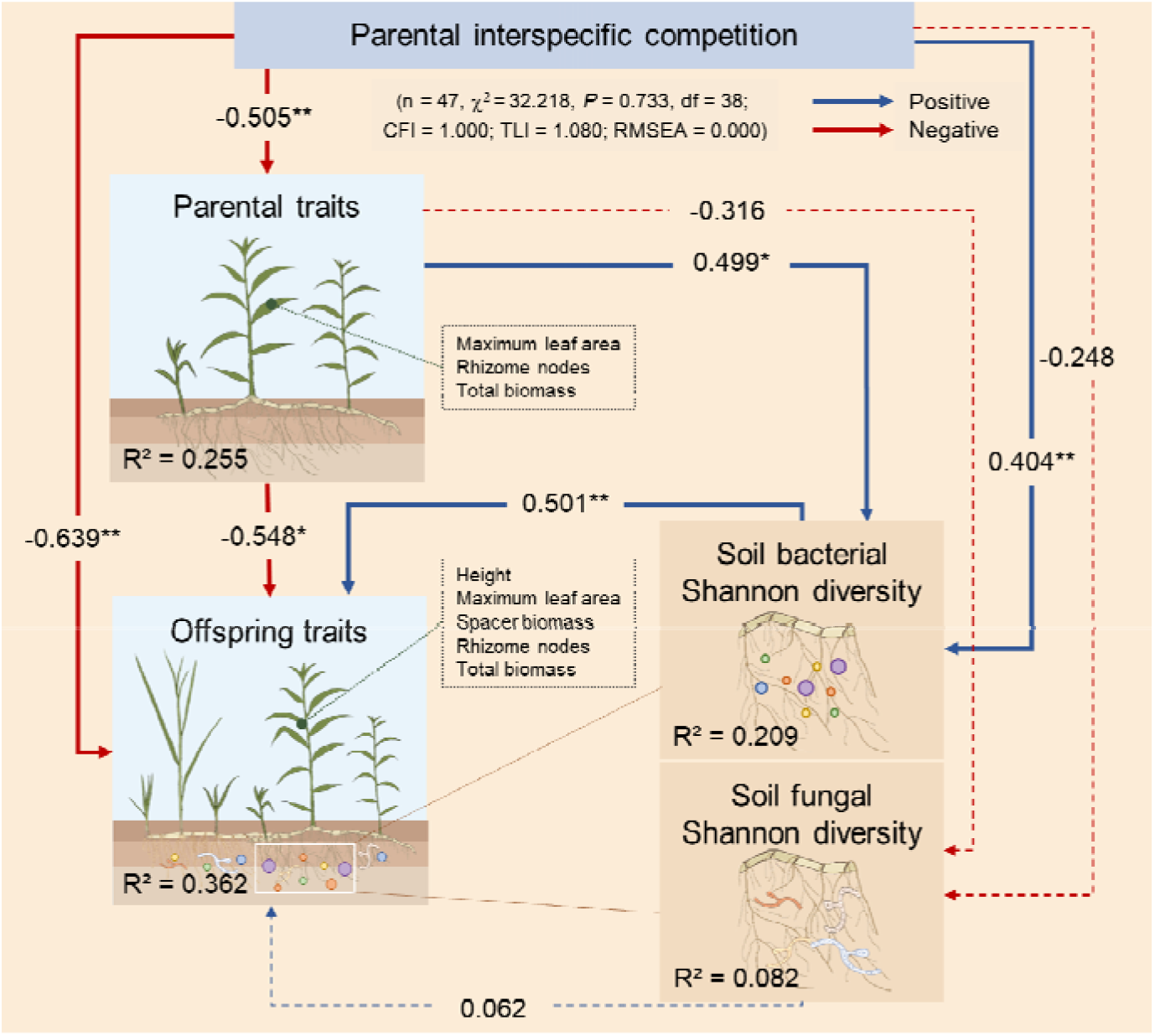
Structural equation model testing direct and indirect relationships between the parental competition environment, phenotypic traits of parent and offspring plants, and the diversity of offspring rhizosphere soil microbes. Blue and red arrows indicate positive and negative relationships, with standardized path coefficients on each. Solid lines indicate significant path coefficients (\**P* < 0.05, \*\**P* < 0.01), and dashed lines non-significant ones. The *R*^2^ values in the boxes indicate the amounts of variance explained by the explanatory variables in each category.

## Discussion

The environmental conditions in which plants grow can affect their offspring through a range of molecular and physiological mechanisms (Liu *et al*., 2022; Sammarco *et al*., 2025). Here we show that even the diversity of plant-associated microbes can be influenced by the competitive environment of the previous plant generation, and that these microbiome changes are related to changes in plant phenotype – a novel aspect of parental effects in plants.

### Parental competition influences offspring phenotypes

It is well-established that parental exposure to abiotic stresses, e.g. salinity, drought, nutrient limitation or light availability, can transmit maternal information to offspring plants (Morgan & Donohue, 2022; Liu *et al*., 2023b; Aycan *et al*., 2024). Such transgenerational cues can influence offspring performances, including germination, photosynthetic capacity, biomass allocation, or reproductive strategies (Zhou *et al*., 2022; Binder *et al*., 2024). Moreover, parental water and nutrient availability can also influence the competition ability of offspring plants (Li *et al*., 2021a; Jin *et al*., 2023; Springer *et al*., 2024).

One question about parental effects that has so far received less attention is whether parental competition experience itself can have transgenerational effects (Puy *et al*., 2021). In our study we found that parental exposure to interspecific competition indeed affected the phenotypes of *P. australis* across generations. Parental competition decreased maximum leaf area, and it also influenced the photosynthetic capacity of the offspring. Previous research often found that plants increase leaf area in response to interspecific competition, presumably to enhance light utilization capacity (Waterman & Sultan, 2021). In our study, however, *P. australis* offspring appeared to prioritize photosynthetic efficiency over leaf growth, possibly because interspecific competition created a low-resource environment for the parent plants (as indicated by their decreased biomass). According to Dong et al. (2018), persistent competition across parent-offspring generations may inhibit offspring growth or enhance competitive ability. Our findings are consistent with these predictions, i.e. decreased leaf area but maintained photosynthetic capacity, and they suggest that transgenerational priming by parental competition may induce a conservative resource-use strategy in clonal offspring of *P. australis*. Interestingly, similar results were found in the apomictic plant *Taraxacum brevicorniculatum* (Puy *et al*., 2021).

### Parental competition alters rhizosphere microbes of offspring plants

Plants often recruit specific soil microbial communities to modify their abiotic or biotic stress resistance, e.g. by exuding metabolites in the rhizosphere (Xu *et al*., 2024; Fu *et al*., 2025). Our study indicates that such environment-dependent plant-microbe associations may also act across generations, as rhizosphere microbial communities were significantly influenced by the parental competition environment. To our knowledge, only two previous studies, on *Tanacetum vulgare* and *Eucalyptus grandis*, respectively (Vivas *et al*., 2017; Malacrinò *et al*., 2025) have demonstrated similar transgenerational effects on offspring microbiome, albeit both focusing on leaf-associated microbes. Here, we demonstrated such effects for the first time for rhizosphere microbes. Specifically, we found that when offspring plants were grown in the same competitive environment as their parents, interspecific competition transgenerationally enriched several offspring-associated bacterial taxa. We can only speculate about the mechanistic underpinnings of these changes, which may be promotion of particular microbes associated with nutrient cycling and stress resistance (Cao *et al*., 2024; Malacrinò *et al*., 2025), possibly influenced through root exudates of the host plants (Kong *et al*., 2019).

Interestingly, we found that relationships between plants and rhizosphere microbes were to some extent genotype-specific, in particular for bacterial communities (Yue *et al*., 2024), and that also the transgenerational effects on root-associated bacteria were influenced by the plant genotype. The differences between genotypes may result from variation in physiological traits, secretion profiles, nutrient demands, or other processes that influence bacteria and thus result in differential recruitment of soil microbial communities (Kudjordjie *et al*., 2021; Przybylska *et al*., 2024; Liu *et al*., 2024). Through their effects on microbial diversity and/or promotion of functionally specialized microbes, these processes may ultimately impact ecosystem stability (Shankar *et al*., 2023; Vincze *et al*., 2024).

### Microbe-mediated parental effects: linking microbes and plant phenotypes

One of the most intriguing results of our study was that the diversity of rhizosphere microbes was related to plant traits both within and across generations. It is known that plant-associated microbial communities are influenced by plant traits (Wang *et al*., 2020), but usually these effects are expected to be direct and within generations. Our mantel tests found that parental traits exerted even stronger effect on offspring bacteria communities than offspring traits, and these results were further supported by the SEM analyses where only parental traits were significantly associated with offspring rhizosphere bacteria while offspring traits were not. The analyses identified several pathways through which parents may have influenced offspring phenotypes:

First, parental competition had a direct negative effect on offspring performance traits, presumably through a reduction in resource provisioning (Gundel *et al*., 2017; Germain *et al*., 2019). Secondly, parental competition negatively impacted parental traits but indirectly positively influenced offspring traits. This may be mediated by parental ‘competition memories’ transmitted through epigenetic cues, thereby promoting offspring pre-adaptation to recurrent competition (Puy *et al*., 2021; Miryeganeh, 2021; Latzel *et al*., 2023). Most importantly, we identified a previously unreported pathway in which parental effects influenced offspring phenotypes through altering offspring rhizosphere microbes. If parental competition increases rhizosphere bacterial diversity across generations, and this bacterial diversity has a positive effect on offspring performance, this may partly compensate for negative effects of parental competition on offspring performance, and the observed phenomenon may be considered a microbial rescue effect.

The precise mechanism underlying these observed microbe-mediated parental effect remains unknown, and we can only speculate. One possibility is that parental competition alters the expression or epigenetic regulation of genes associated with synthesis of root exudates which could contribute to heritable differences in offspring microbe recruitment (Yan *et al*., 2024; Yaghoubi Khanghahi *et al*., 2024; Gopalan-Nair *et al*., 2024; Liu *et al*., 2026). Another possibility is that parental competition alters physiological signals transmitted to clonal offspring such as changes in secondary metabolites, phytohormones, antioxidants, or other signaling compounds (Kong *et al*., 2019; Jacoby *et al*., 2021; Kudjordjie *et al*., 2021; Bastías *et al*., 2022; Liu *et al*., 2024). These signals might subsequently influence offspring root exudates and promote the recruitment of specific plant growth-promoting bacteria, thereby restructuring rhizosphere microbial communities (Trivedi *et al*., 2022; Zhao *et al*., 2024; Malacrinò *et al*., 2025; Johnson & Marín, 2025). This is consistent with previous evidence of rapid responses of plant-associated microbes to environmental changes and their cross-generational effects on plant performance (Varga *et al*., 2013; Vivas *et al*., 2015; Gundel *et al*., 2017; Suryanarayanan & Shaanker, 2021; Fort *et al*., 2021; Anand *et al*., 2023; Latzel *et al*., 2024; Beltrán-Torres *et al*., 2026). Unlike mechanisms based on vertical transmission of endophytic microbes or classical plant-soil feedback (PSF) (Bennett & Klironomos, 2019), our results suggest that parental plants may influence offspring rhizosphere microbial assembly even when offspring are grown in fresh soil. Thus, parental competition may establish a form of transgenerational microbial recruitment that benefits offspring under predictable competitive conditions.

If microbes are the mediators of parental effects, then our results indicate that it is bacteria rather than fungi that are involved in this. This may be because bacteria respond more rapidly to environmental stimuli, whereas many fungi, in particular symbiotic species, may depend more strongly on stable host associations and may be more susceptible to stochastic environmental variation (Powell *et al*., 2015; Hassani *et al*., 2018; Li *et al*., 2024).

Overall, our results extend the current understanding of parental effects by demonstrating that parental environmental conditions can influence offspring phenotypes not only directly, but also through alteration of rhizosphere microorganisms.

## Conclusions

Here we found a new pathway through which offspring rhizosphere microbial diversity mediates parental effects on offspring phenotypes. Parental effects could shape offspring performance through plant-microbe interactions processes, offering a more holistic perspective on plant adaptation to the predictable competition. Although our study provides evidence for the role of horizontal rhizosphere microbial recruitment in regulating plant transgenerational plasticity, the specific mechanisms and signaling pathways still require further in-depth investigation. We expect future studies to reveal how parental effects work together in genetic, epigenetic and ecosystem dynamics to shape adaptive evolutionary trajectories in plants.

## Acknowledgements

This work was supported by the National Natural Science Foundation of China (grant numbers 32371584, 32071525 to FLL, 32602177 to YHC) and a China Scholarship Council grant (Project 202306510039) to YHC, as well as the Deutsche Forschungsgemeinschaft (projects BO 3241/11-1 and BO 3241/10-1 to OB) as part of the DFG priority program 2125 Deconstruction and Reconstruction of the Plant Microbiota “DECRyPT”. We thank the High Performance and Cloud Computing Group at the Zentrum für Datenverarbeitung of the University of Tübingen for managing the BinAC server.

## Competing interests

The authors declare no competing interests.

## Author contributions

FLL and YHC designed the study. YHC, CLW, TF, YFL and JQM performed the experiments. YHC, JHJ, and FR analyzed the data, with input from MP and OB. YHC wrote the first draft of the manuscript. All authors contributed to revision and improvement of the manuscript.

## Notes

### Competing Interest Statement

The authors have declared no competing interest.

