## Supporting Information for "Parental effects can be transmitted through the rhizosphere microbiome"

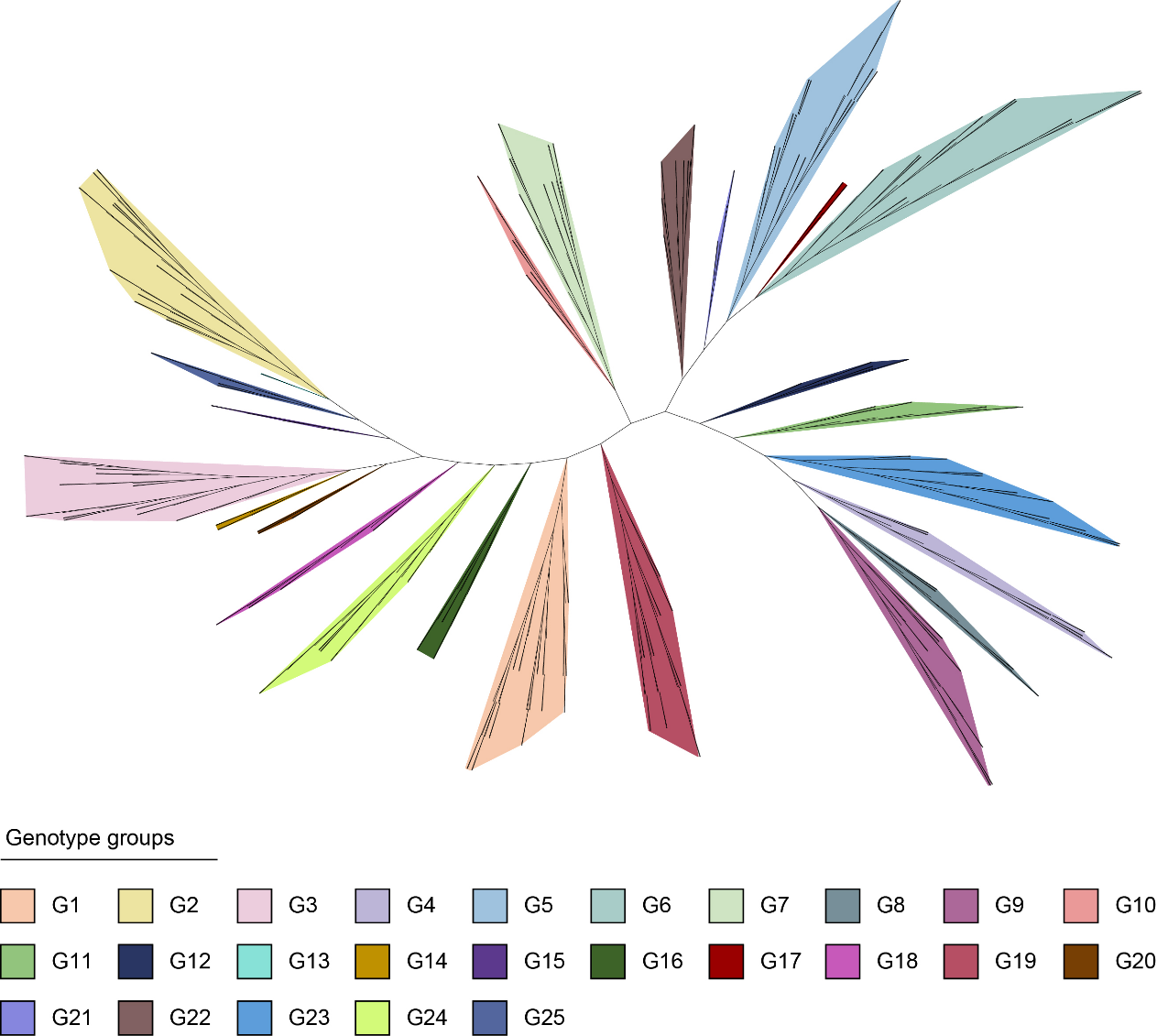

### Fig. S1 Genetic relationships, based on previously published Simple Sequence Repeat marker data [45], among 416 *Phragmites australis* individuals from 42 populations in four estuarine wetlands (Liaohe Estuary, Yellow River Estuary, Yangtze Estuary, and Min River Estuary) along the Chinese coast. The phylogenetic tree is based on Nei's genetic distances, using the neighbor-joining method.

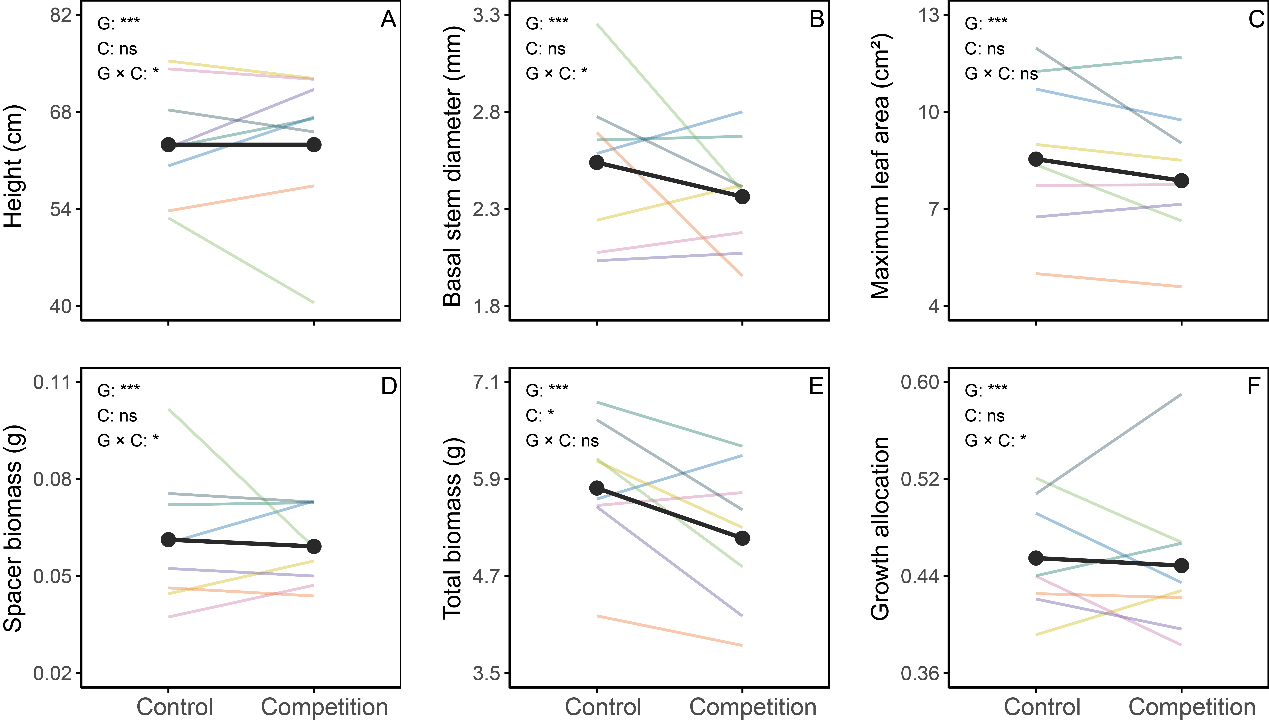

### Fig. S2 Direct effects of interspecific competition on plant phenotypes (parental generation): (A) plant height; (B) basal stem diameter; (C) maximum leaf area; (D) spacer biomass; (E) total biomass; and (F) growth allocation. The coloured lines represent different genotypes, and the thick black lines are the respective main effects (across genotypes). All values are predicted values from linear mixed-effects models, back-transformed to the original scale. The significances of fixed effects are indicated in the figure panels, with *** *P* < 0.001, * *P* < 0.05 and ns *P* > 0.05 (G: genotype, C: competition).

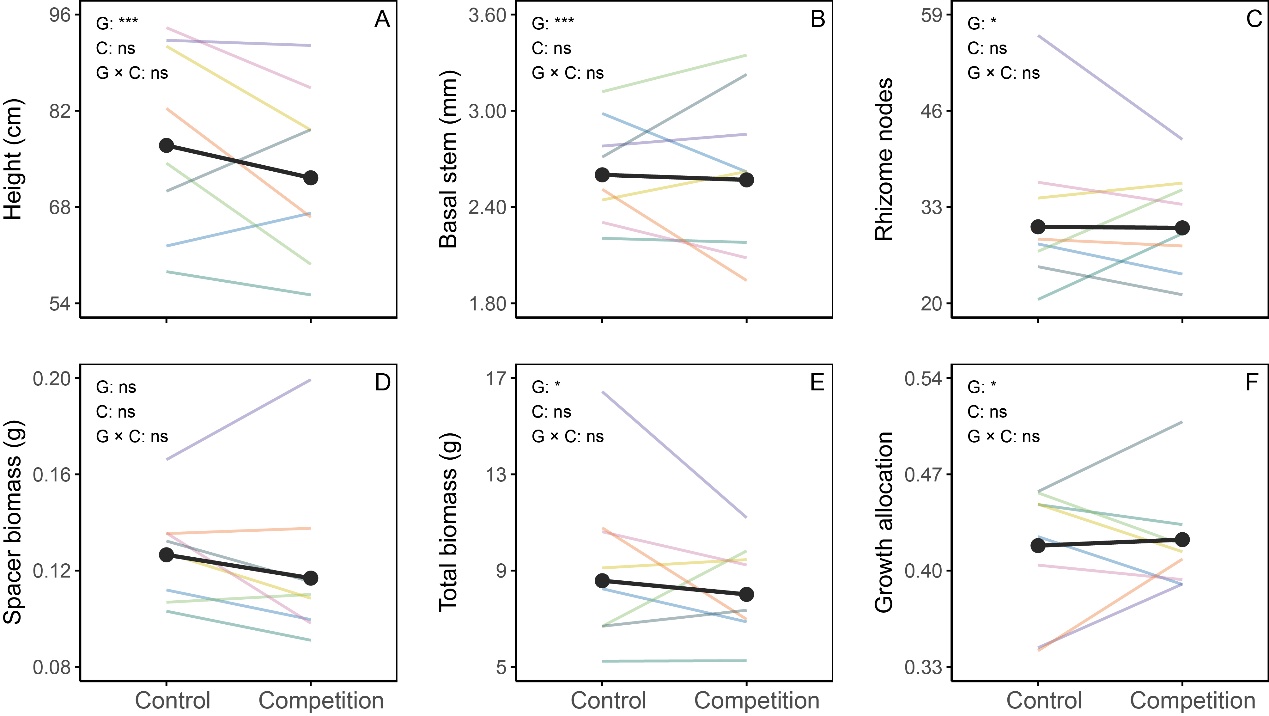

### Fig. S3 Effects of parental competition environment on phenotypic traits of offspring plants in *P. australis*: (A) plant height; (B) basal stem diameter; (C) the number of rhizome nodes; (D) spacer biomass; (E) total biomass; and (F) growth allocation. The coloured lines represent different genotypes, and the thick black line is the main effect (across genotypes) of parental competition. All values are predicted values from linear mixed-effects models, back-transformed to the original scale. The significances of fixed effects are indicated in the figure panels, with *** *P* < 0.001, * *P* < 0.05 and ns *P* > 0.05 (G: genotype, C: parental competition).

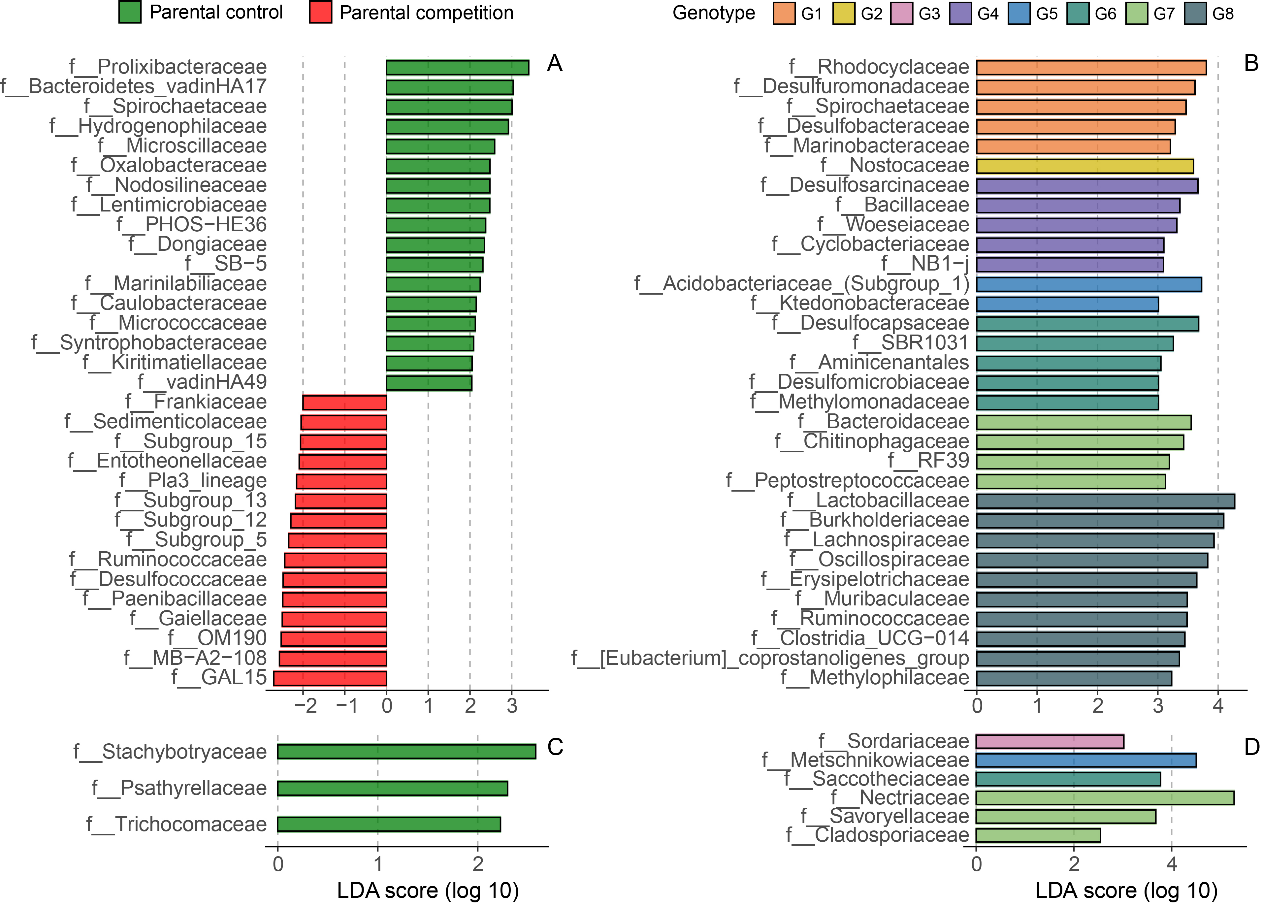

### Fig. S4 Microbial taxa significantly associated with parental competition treatments (A: bacteria, C: fungi) or specific *P. australis* genotypes (B: bacteria, D: fungi), as identified by linear discriminant analysis effect size (LEfSe) analyses. We used a linear discriminant analysis (LDA) score threshold of >3 for bacteria in the genotype comparison (panel B), and a threshold of > 2 for all other tests.
